# Knowledge, attitudes and practices of hypertension in a community based cross sectional study done in Ward 14, Gwanda District, Matebeleland South, Zimbabwe

**DOI:** 10.1101/599688

**Authors:** Pugie Tawanda Chimberengwa, Mergan Naidoo

## Abstract

**Background:** Hypertension is a significant contributor to cardiovascular and renal diseases. In poor communities there is lack of awareness, poor treatment and control. However, it can be controlled by lifestyle modifications. The aim of this study was to determine knowledge, attitudes and practices with regards to hypertension in a rural disadvantaged community in Matebeleland South province of Zimbabwe.

**Methods:** We conducted a descriptive cross-sectional survey. A pre-tested and validated interviewer administered questionnaire was used to collect demographic, awareness, treatment and control data among consenting hypertensive patients.

**Results:** 304 respondents were enrolled into the study, their mean age was 59 years and 65.4% were females. Adding salt on the table (59.8%) was a risk factor. There were strong community beliefs in managing hypertension with herbs (50.7%) and use of traditional medicines (14.5%). Knowledge on hypertension was poor with 43.8% of hypertensive patients having had a discussion with a health worker on hypertension and 64.8% believing the main case of hypertension is stress while 85.9% stated palpitations as a symptom of hypertension. Defaulter rate was high at 30.9% with 25% of those on medication not knowing whether their blood pressure control status. Odds ratio for good knowledge for secondary and tertiary education were 3.68 (95%CI: 1.61-8.41) and 7.52 (95%CI: 2.76-20.46) respectively compared to no formal education. Those that believed in herbal medicines and those that used traditional medicines were 53% (95%CI: 0.29-0.76) and 68% (95%CI: 0.29-0.76) less likely to have good knowledge compared to those who did not believe and use traditional medicines respectively.

**Conclusion:** Lack of education and poor socio-economic backgrounds were associated with poor knowledge on hypertension. Shortages of medication, poor health funding and weak health education platforms contributed to reduced awareness and control of hypertension in the community. Thus, community hypertension awareness, treatment and control needed to be upscaled.

## Introduction

Hypertension (HT) is one of significant health challenges in low- and middle-income countries (LMICs) that are experiencing epidemiological transition from communicable to non-communicable diseases [1-3]. Hypertension is prevalent in both high income and LMICs. Together with other cardiovascular diseases, these public health problems that are strongly linked to urbanization, aging populations, westernized socio-economic sedentary lifestyles promoting excessive salt and alcohol intake, smoking, obesity as well as lack of physical exercise [4-7].

Hypertension is the most common incidentally diagnosed chronic disease and various factors affect diagnosis, treatment and control of HT. However, the most important barrier to diagnosis is lack of knowledge and awareness on HT and its complications [8]. Almost half of hypertension-related deaths can be averted with compliance or adherence to antihypertensive treatment [9]. It is therefore important to assess the patients’ knowledge and awareness on HT because patient education is a key component in the programs and interventions designed to control HT [10].

Hypertension is a major risk factor for cerebro-vascular accidents as well as coronary heart diseases, with two-thirds of all cerebro-vascular accidents being attributable to poor HT control [11,12]. Cardiovascular diseases are the major cause of death globally, with an estimated 17.5 million deaths per year and 80% of the deaths are recorded in LMIC [5,9]. In African communities, the challenges in managing HT lie in prevention, diagnosis and treatment [13]. There is a shortage of national data on HT prevalence studies in Zimbabwe. A study that summarized HT prevalence over a 14-year period from 1997 to 2010 estimated the pooled prevalence of HT in Zimbabwe at 30% [14]. In a hypertension study done in Bulawayo city, in southern Zimbabwe, the highest prevalence of 38.4% was reported [15] while an average prevalence of 17.9% was recorded among three provinces in another survey focusing on both urban and rural settings [16].

Significant progress has been made in improving HT awareness, treatment and control among patients living with hypertension (PLWHT). Efforts to control HT have included improving public knowledge and awareness on the risks and complications of hypertension [10]. The aim of this study was to determine the level of knowledge, attitudes and practices with regards to HT in ward 14, Gwanda district (Zimbabwe) as part of a community based participatory research (CBPR) we carried out. This paper will report on the quantitative baseline findings on the community’s knowledge and awareness on hypertension.

## Methods

### Study design

We conducted a baseline descriptive cross-sectional study to evaluate knowledge, awareness and perceived control of hypertension.

### Study setting

The survey was undertaken in Ward 14, a rural area situated about 50km south-west of Gwanda town, Matebeleland South province in Zimbabwe. Gwanda district population was 115 778 inhabitants which was 16,9% of the provincial population. Ward 14 had 1384 households, 5867 inhabitants of which 55% were females [20].

### Sampling and Sample size calculations

Using the Dobson formula; n =(z^2^pq)/Δ^2^, where n = sample size, z = standard error risk, p = prevalence of hypertension (PLWHT), q = 1-p (proportion of people without hypertension) and Δ = absolute precision. Assuming 95% CI (z=1.96), a prevalence of HT (p) of 27% [16], and using a precision of 5%, it was established that an adequate sample size would comprise 303 PLWHT. All consenting hypertensive patients from ward 14 drawn from Sengezane rural health center, those identified in the villages, schools and business premises and hypertensive patients that were seen at Gwanda provincial hospital (primary referral center for Sengezane clinic) whose address originated from ward 14 were enrolled into the study.

### Ethics approval and consent to participate

All phases of this research were jointly approved by the Medical Research Council of Zimbabwe (MRCZ/A/2136) and the Biomedical Research Council of the University of Kwa-Zulu Natal, South Africa (BFC318/16). Authority was sought from the Ministry of Health and Child Care Zimbabwe through the Provincial Medical Director, Matebeleland South and the District Medical Officer for Gwanda. Gatekeeper’s authority was sought from the Ministry of Local Government at the offices of District Administrator. The Chiefs, headmen, religious leaders, health center committee and community advisory board were approached through Ward 14 councilor. Written informed consent was sought from all research participants.

### Data collection

Quantitative data was collected using interviewer administered questionnaires. All self-reported persons to have been diagnosed with HT in the ward qualified to be enrolled into the study. The inclusion criteria were any consenting person above the age of 18 years residing in ward 14 who reported having been diagnosed of HT regardless of whether they were taking anti-hypertensive medication or not. The exclusion criteria were any person who was hypertensive not consenting to the study and those below the age of 18 years.

Data collection was done by cooperative inquiry group (CIG) members. These comprised of hypertensive patients, community leaders and village health workers who were joined by nurses and the principal investigator (PI) to form the CIG. They were trained in quantitative data collection using interviewer administered questionnaires on HT knowledge, awareness, treatment and control, recording of data for standardization and uniformity including ethical issues. The PI developed the interviewer administered questionnaire which was discussed, validated and adopted by the CIG for data collection. The validation of the tool was done during a CIG meeting where the participants discussed and understood the meaning of each of the questions in the data collection tool, recording and interpretation of the responses. Each CIG member was given five copies of the questionnaire to use as a pre-test and responses were shared and agreed upon. Thereafter, a full data collection exercise over a period of 2 weeks was then commissioned. Data collected included respondents’ demography, risk factors for hypertension, knowledge, attitudes, perceptions and barriers to treatment compliance.

### Statistical methods

Frequencies and proportions for lifestyle related factors and knowledge were calculated as well as specific proportions for age, marital status, income and level of education. The proportion of hypertensive patients who were aware of their diagnosis, receiving treatment, BP control was determined including time of last BP measurement. The proportion of hypertensive patients on treatment to those aware of their blood pressure control was assessed. Ten key questions on hypertension knowledge and practice were recoded and good knowledge was ascertained by getting at least six questions correct respectively while getting more than three correct questions was regarded as good attitude. This was used for bivariate logistic regression analysis to analyze factors influencing hypertension. Data was analyzed using Microsoft excel, Stata and Epi-Info 7 software.

### Study validity and bias

Selection bias was reduced in the study by ensuring high participation rates. We enrolled almost all possible hypertensive patients in the ward to approximate the whole population of PLWHT. Interviewer bias was reduced by using trained CIG members who had standardized questionnaires. To reduce reporting bias, the respondents were asked for the signs and symptoms of HT they knew instead of selecting from a list. The participation of GIG members drawn from the community for data collection while embedding questions on specific lifestyles and putting less emphasis on non-desirable social habits reduced social desirability and information bias. In instances, where clinic records were available, they were utilized for triangulation and to reduce information bias.

## Results

A total of 304 PLWHT were enrolled into and participated in the study and the mean age of the participants was 59 (Q1-Q3; 46-72) years. Table 1 shows the sociodemographic data for hypertensive patients who were enrolled into the study.

**Table 1;.**
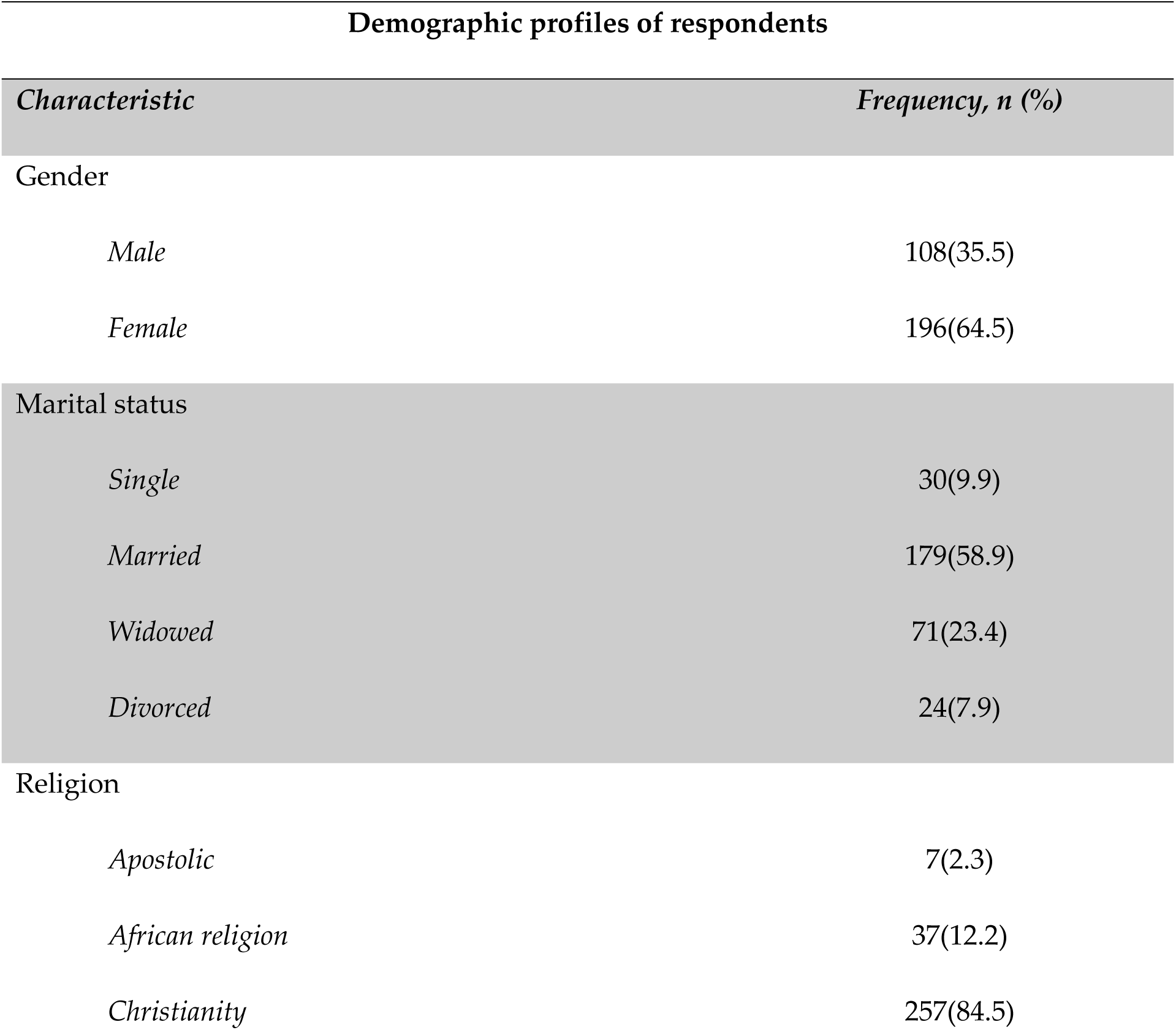
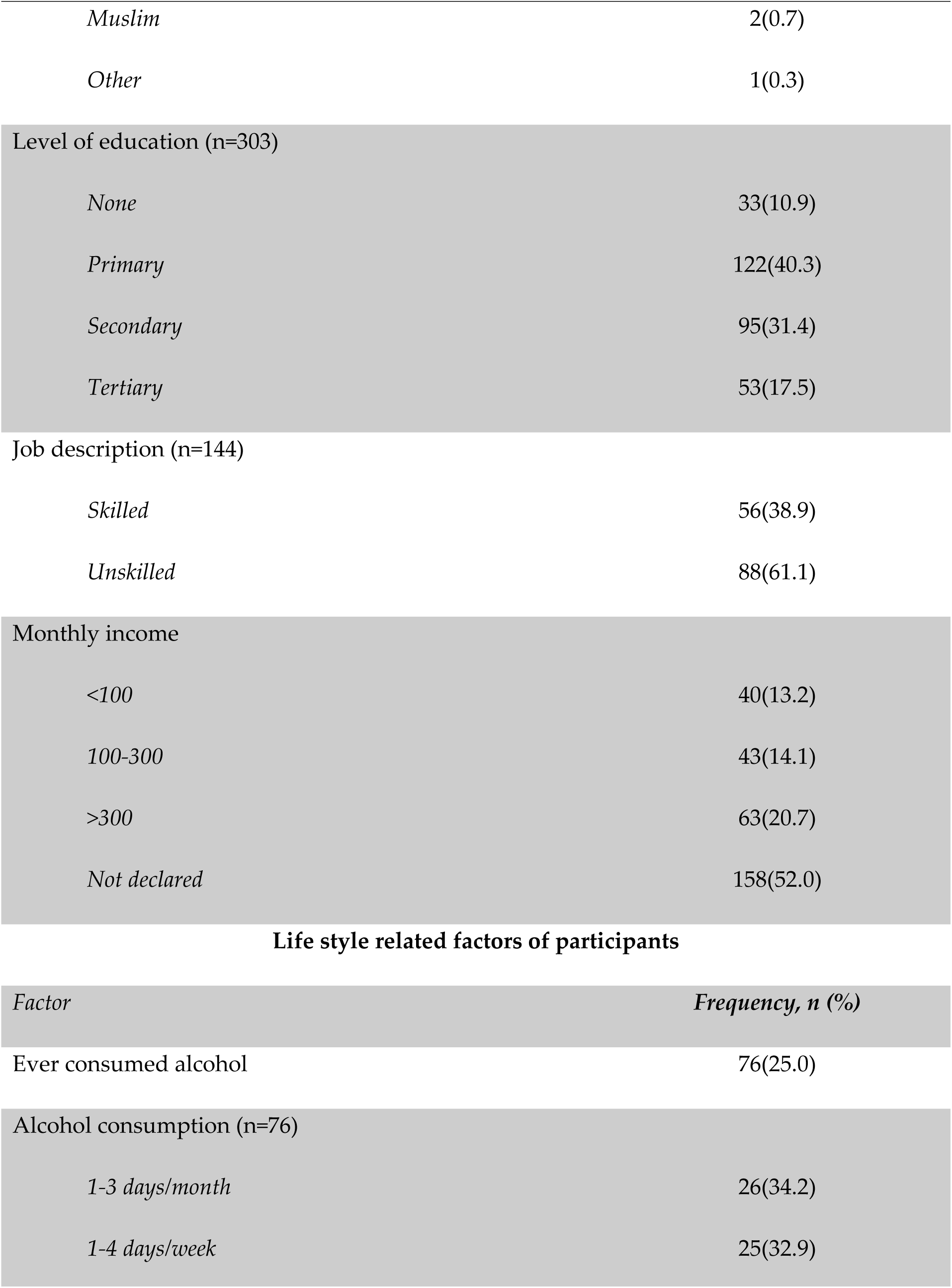
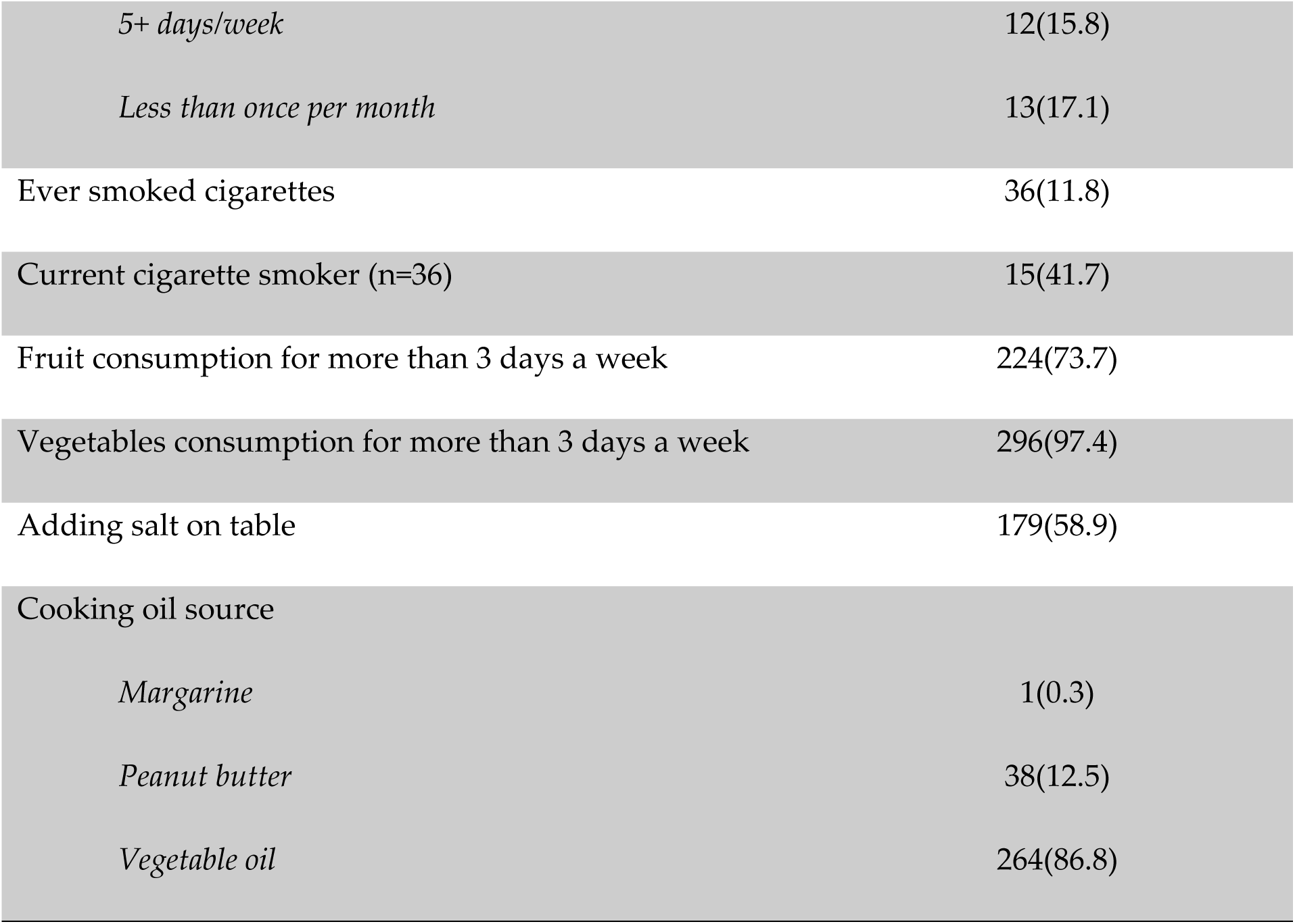
Socio-demographic data and lifestyle related factors for PLWHT; ward 14, Gwanda district; community-based action research project, May 2017.

| Demographic profiles of respondents |  |
| --- | --- |
| *Characteristic* | *Frequency, n (%)* |
| Gender |  |
| *Male* | 108(35.5) |
| *Female* | 196(64.5) |
| Marital status |  |
| *Single* | 30(9.9) |
| *Married* | 179(58.9) |
| *Widowed* | 71(23.4) |
| *Divorced* | 24(7.9) |
| Religion |  |
| *Apostolic* | 7(2.3) |
| *African religion* | 37(12.2) |
| *Christianity* | 257(84.5) |
| *Muslim* | 2(0.7) |
| *Other* | 1(0.3) |
| Level of education (n=303) |  |
| *None* | 33(10.9) |
| *Primary* | 122(40.3) |
| *Secondary* | 95(31.4) |
| *Tertiary* | 53(17.5) |
| Job description (n=144) |  |
| *Skilled* | 56(38.9) |
| *Unskilled* | 88(61.1) |
| Monthly income |  |
| *<100* | 40(13.2) |
| *100-300* | 43(14.1) |
| *>300* | 63(20.7) |
| *Not declared* | 158(52.0) |

Life style related factors of participants
| *Factor* | *Frequency, n (%)* |
| --- | --- |
| Ever consumed alcohol | 76(25.0) |
| Alcohol consumption (n=76) |  |
| *1-3 days/month* | 26(34.2) |
| *1-4 days/week* | 25(32.9) |
| *5+ days/week* | 12(15.8) |
| *Less than once per month* | 13(17.1) |
| Ever smoked cigarettes | 36(11.8) |
| Current cigarette smoker (n=36) | 15(41.7) |
| Fruit consumption for more than 3 days a week | 224(73.7) |
| Vegetables consumption for more than 3 days a week | 296(97.4) |
| Adding salt on table | 179(58.9) |
| Cooking oil source |  |
| *Margarine* | 1(0.3) |
| *Peanut butter* | 38(12.5) |
| *Vegetable oil* | 264(86.8) |

The study sample consisted of 196 (64.5%) females and 108 (35.5%) men. Most respondents (47%) were above the age of 60 while only 3% were below the age of 30. The respondents were predominantly Christians 84.5% while the African tradition was observed by 12.2%. In terms of formal education, 51.2% attempted primary level or below with about 11% having not attended primary school at all. Fifty two percent of the respondents did not declare their family monthly income with only 21% declaring monthly family income above US$ 300. As shown in Table 1, 75% of respondents did not drink alcohol with only 25% reported to have used alcohol mainly as social drinkers and 12% had ever smoked. Twenty four percent ate fruits on more three days per week, 67% ate vegetables on more than three days a week while 60% reportedly added salt on the table. Vegetable oil was used for cooking by 87% of respondents.

### Knowledge and beliefs on HT

Table 2 shows the respondents’ knowledge and beliefs on HT treatment and control among respondents.

**Table 2;.** Knowledge and beliefs on hypertension, treatment and control among HT patients in ward 14, Gwanda district community-based action research project on hypertension, May 2017.

| Knowledge category | Frequency, n (%) |
| --- | --- |
| Positive family history of hypertension | 204(67.1) |
| Positive family history of known hypertension complications | 116(39.3) |
| Belief in effectiveness blood pressure pills to manage HT | 285(93.8) |
| Belief in herbal remedies to manage HT | 153(50.7) |
| Agree to using African medicines to manage HT | 44(14.5) |
| Can use traditional medicines to control your blood pressure | 44(14.5) |
| Source of knowledge on high blood pressure |  |
| *Local clinic nurse* | 172(56.6) |
| *Village health worker* | 35(11.5) |
| *Public hospital* | 55(18.1) |
| *Private doctor* | 33(10.9) |
| *Other* | 9(3.0) |
| Has a health worker / village health worker ever discussed<br>with you about blood pressure treatment and control? | 133(43.8) |
| What are the causes of blood pressure? |  |
| *Drugs* | 16(5.3) |
| *Old age* | 23(7.6) |
| *Stress* | 197(64.8) |
| *Witchcraft* | 18(5.9) |
| *Unknown* | 48(145.8) |
| *Other* | 2(0.7) |
| What are the signs and symptoms of high blood pressure? |  |
| *Asymptomatic* | 22(7.2) |
| *Headache* | 1(0.3) |
| *Palpitations* | 261(85.9) |
| *Other* | 8(2.6) |
| Don't know | 12(4.0) |
| Can one have high blood pressure without any signs and symptoms? | 161(53.0) |
| What are the consequences of untreated blood pressure? |  |
| *Stroke, Heart failure, Kidney failure* | 278(91.4) |
| *Other* | 7(2.3) |
| *Don't know* | 19(6.3) |
| What are the risk factors for developing high blood pressure? |  |
| *Hereditary* | 10(3.3) |
| *Diet* | 253(83.2) |
| *Don't know* | 41(13.5) |

Family history of HT was reported by 67% of respondents while 39% agreed to knowing a hypertensive family member who developed complications of HT. Ninety four percent believed tablets lower blood pressure and 51% also believed in the use of traditional remedies to lower blood pressure while 15% confessed to visiting traditional healers for blood pressure treatment. The local clinic nurse was primarily the source of knowledge on HT for 57% of the respondents while the VHW was consulted by only 12%. Forty four percent reported having discussed blood pressure treatment control with a health worker or a VHW.

Stress was reportedly cited as the commonest cause (64.8%) of HT and 16.8% reported that HT causes were unknown. Palpitations were thought to be the commonest symptom of HT by 85.9% of respondents while only 7.2% stated that HT was asymptomatic and 4% did not know. Fifty three percent of respondents agreed that one can have HT without presenting any signs and symptoms. Death, stroke, heart and kidney failure were cited as the commonest complications of poorly controlled or untreated HT. Diet was singled out as the commonest risk factor for HT (83.2%) with heredity being mentioned by 3.3% and 13.5% of respondent could not state any risk factor for HT.

### Attitudes and practices on HT

Table 3 shows the attitudes and practices of respondents on the control of HT, their preferred service providers and access areas for follow up.

**Table 3;.** Attitudes and practices on treatment and control of HT in ward 14, Gwanda district community-based action research project on hypertension, May 2017.

| **Treatment and control** | **Frequency, n (%)** |
| --- | --- |
| How do you currently control your blood pressure? |  |
| *Blood pressure tablets* | 260(85.5) |
| *Traditional medicines* | 27(8.8) |
| *Prayer* | 9(3.0) |
| *Other* | 8(2.6) |
| those currently taking medication for hypertension | 250(82.2) |
| Taken medication regularly in the past 2 weeks (n=250) | 229(91.6) |
| Those having challenges with taking hypertensive treatment | 58(19.1) |
| Preferred place of followed up for hypertension management |  |
| *Local clinic* | 159(52.3) |
| *Public hospital* | 48(15.8) |
| *Private doctor* | 49(16.1) |
| *My home* | 13(4.3) |
| *Other* | 3(1.0) |
| *Not stated* | 32(10.5) |
| When last was your blood pressure checked? |  |
| *< 1 month ago* | 152(50.0) |
| *2-4 months ago* | 63(20.7) |
| > 4 months ago | 58(19.1) |
| Missing | 31(10.2) |
| Is your blood pressure well controlled? |  |
| Yes | 196(64.5) |
| No | 32(10.5) |
| Don't know | 46(15.1) |
| Not stated | 30(9.9) |
| Ever defaulted hypertensive medication | 94(30.9) |
| Reasons for defaulting treatment (n=93) |  |
| Had developed side effects from medicines | 11(11.8) |
| Tablets make me sick | 7(7.5) |
| To avoid addiction | 11(11.8) |
| Treatment was not effective | 14(15.1) |
| Trying alternative remedies | 16(17.2) |
| Feeling much better | 26(28.0) |
| Other | 8(8.6) |
| Agreed to use of traditional medicines to control blood pressure? | 44(14.5) |

The majority (85.5%) reported that they were using BP tablets to control their HT while 8.8% used African traditional medicines. Eighty two percent respondents reported that they were currently taking the tablets and of these, 91.6% said they had been swallowing the tablets in the preceding 2 weeks while 19.1% cited facing challenges with compliance to treatment during the same time period. The local clinic (52.3%), private doctor (16.1%) and the public hospital (15.8%) were the places where respondents were being followed up for continued care. Fifty percent reported they had a BP reading checked within the preceding month while 10.2% did not know when they last checked and 19.1% had checked more than 4 months prior. With regards to blood pressure control, 64.5% stated that they had well controlled blood pressure, 10.5% knew they had poorly controlled while 25% did not know and 31% had ever defaulted treatment. Feeling much better (27.6%), assuming treatment was not effective (14.9%), avoiding addiction (11.7%) and experiencing side effects of HT medication (11.7%) were some of the commonest excuses given for defaulting treatment. A significant 14% agreed that they can rely on traditional medicines to control blood pressure.

### Regression analysis

Table 4 shows a logistic regression analysis of factors affecting knowledge on hypertension. Data was recoded such that those who had scored six or more out of ten on knowledge and practice scores were deemed to have good knowledge and good practice respectively whilst, a score of three or more out of five was deemed good attitude towards HT treatment and control.

**Table 4;.**
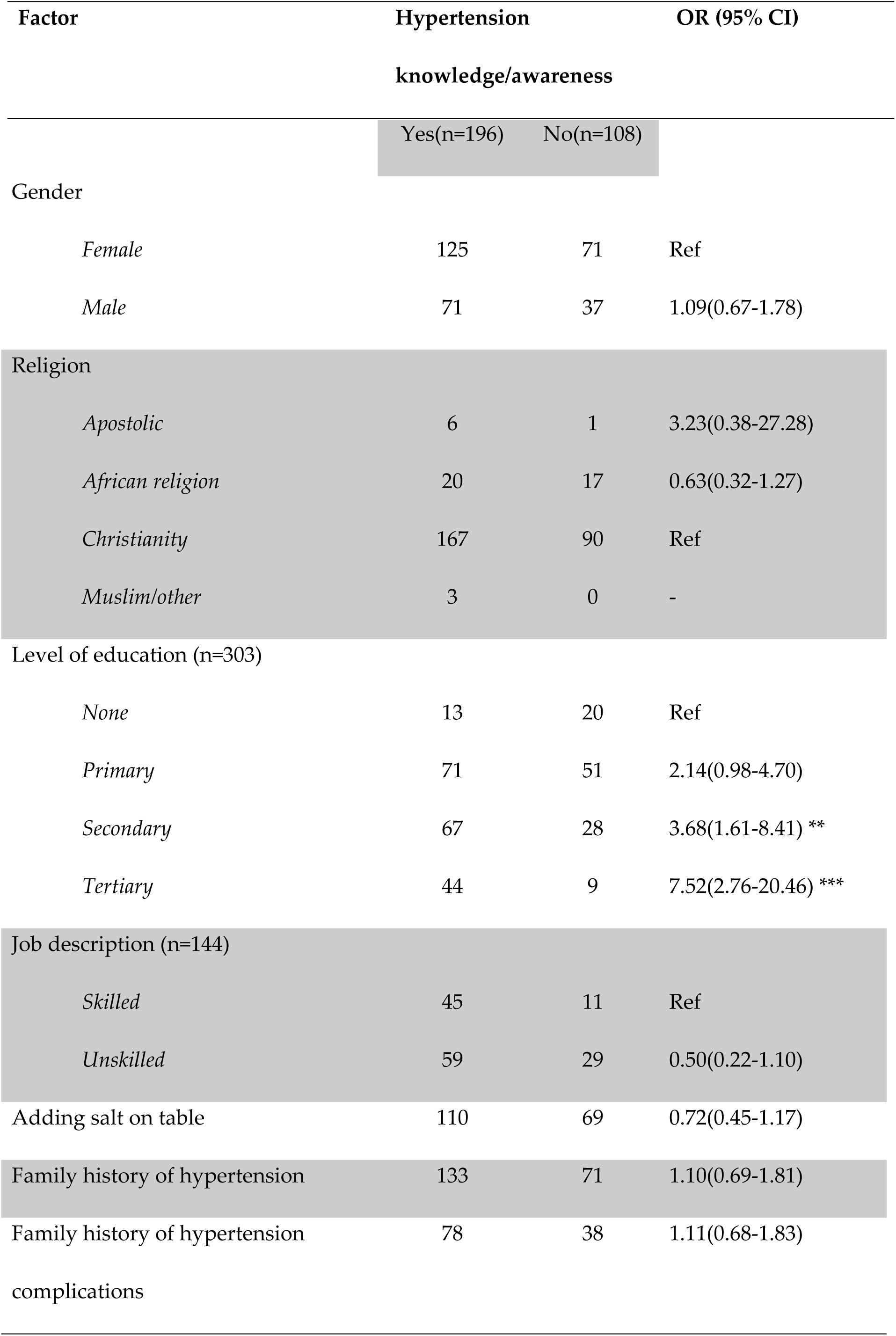
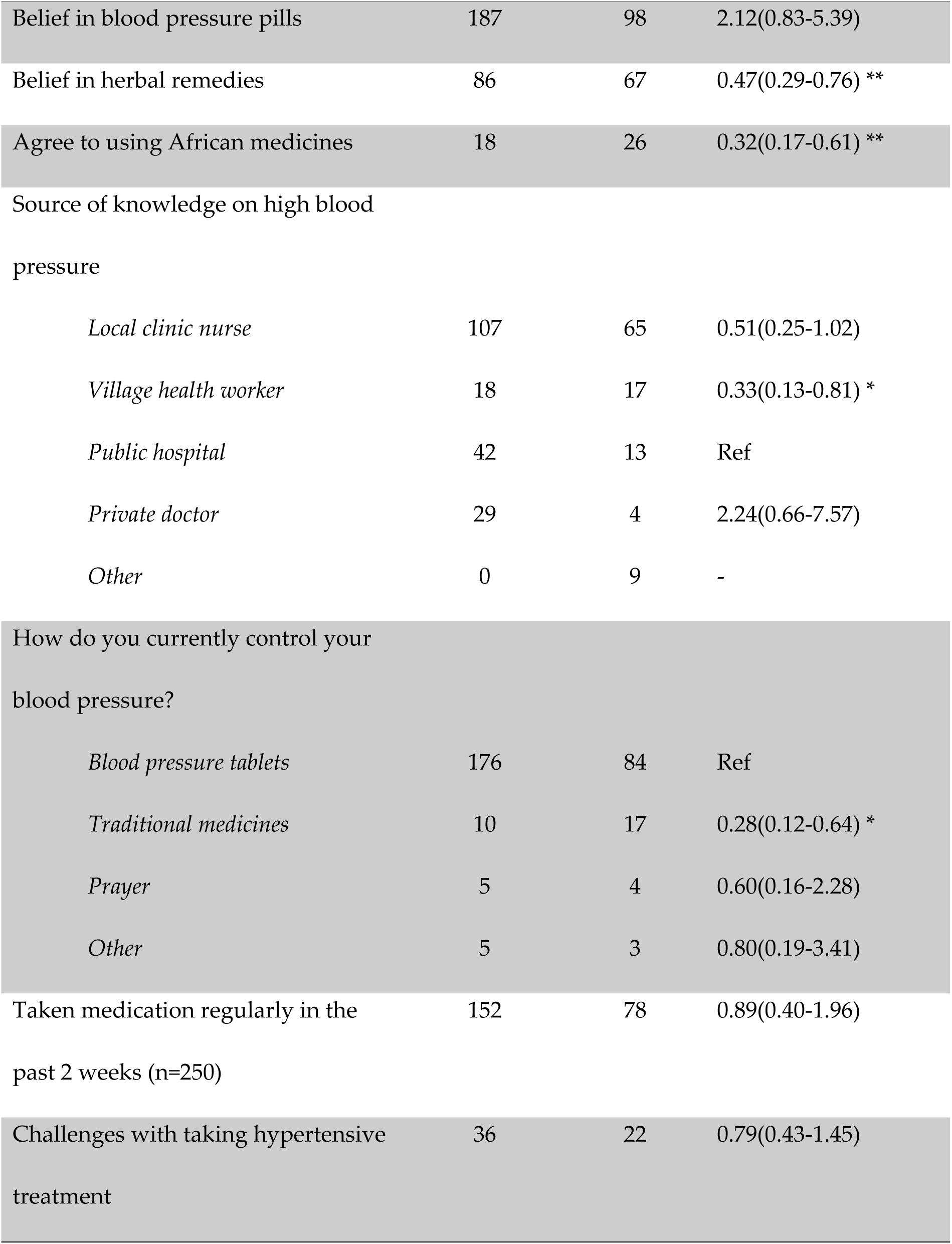
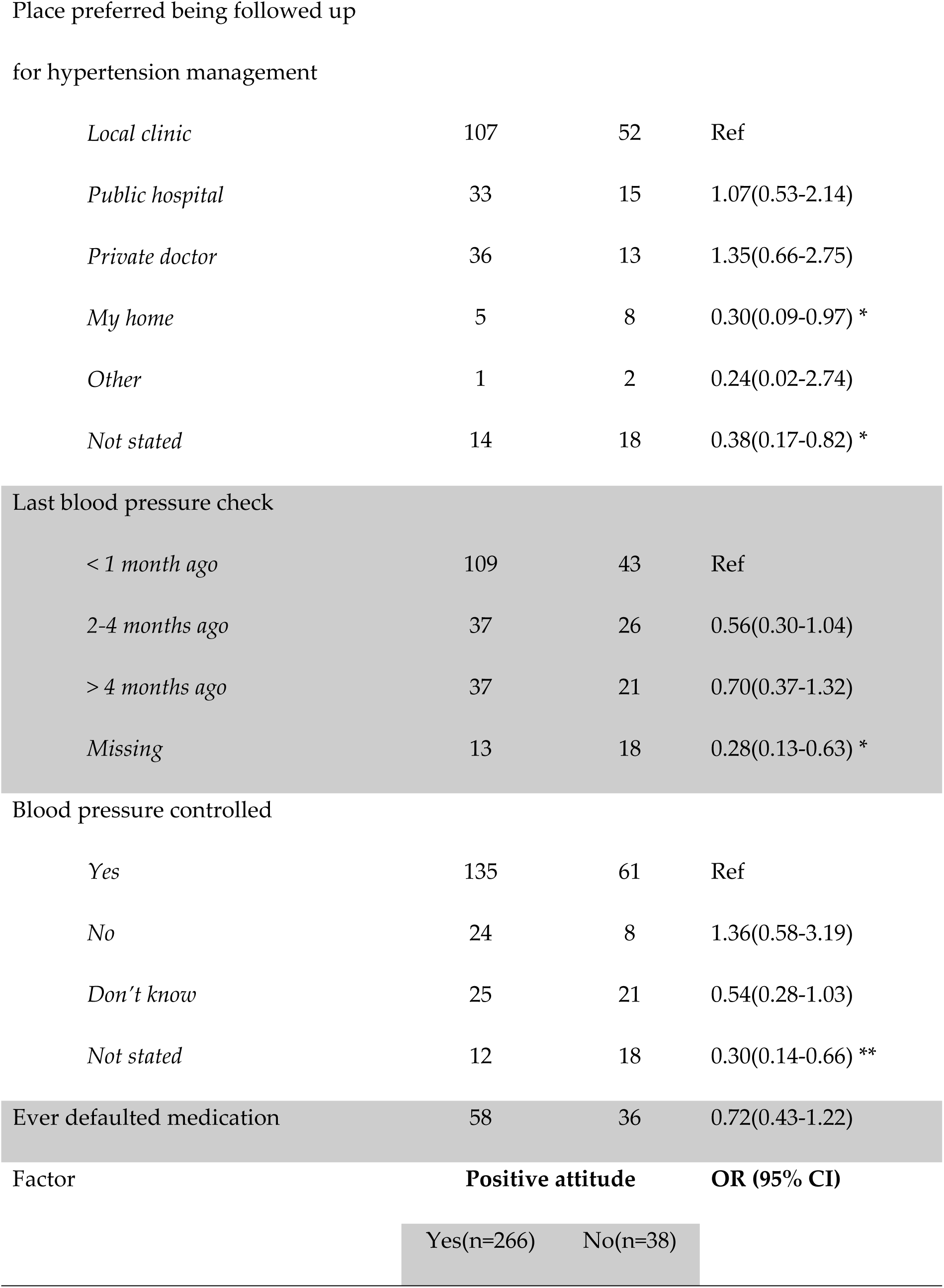
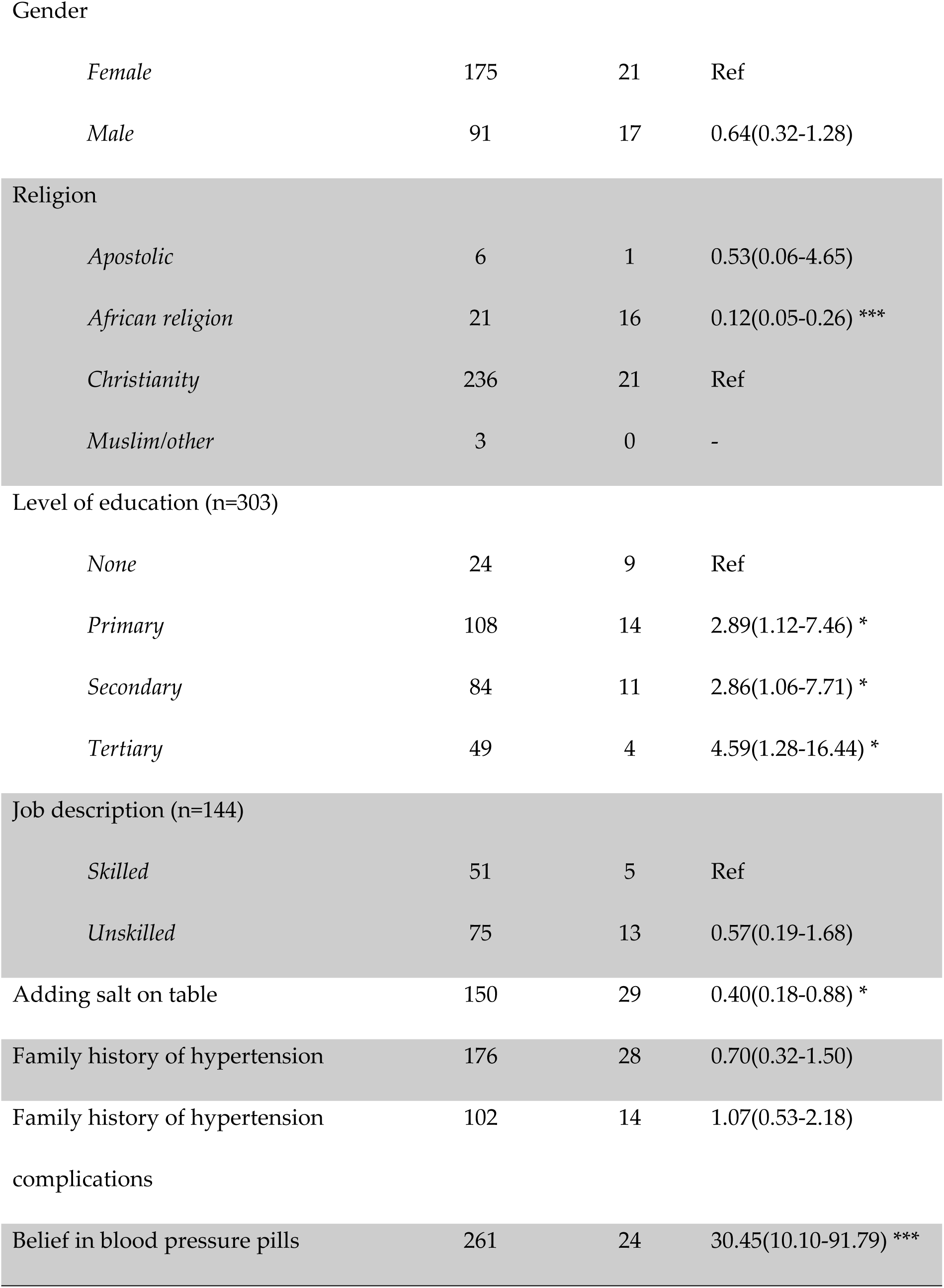
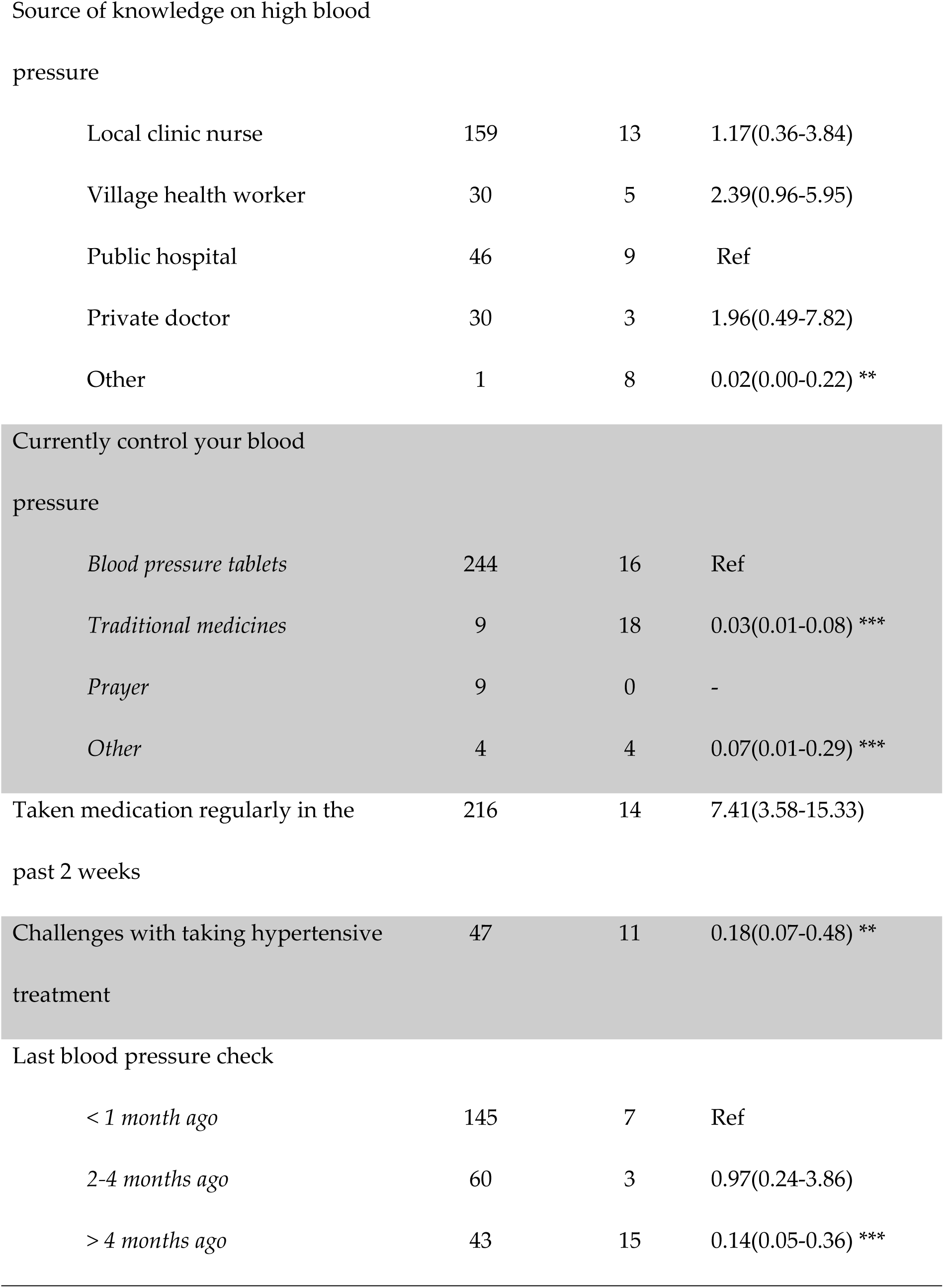
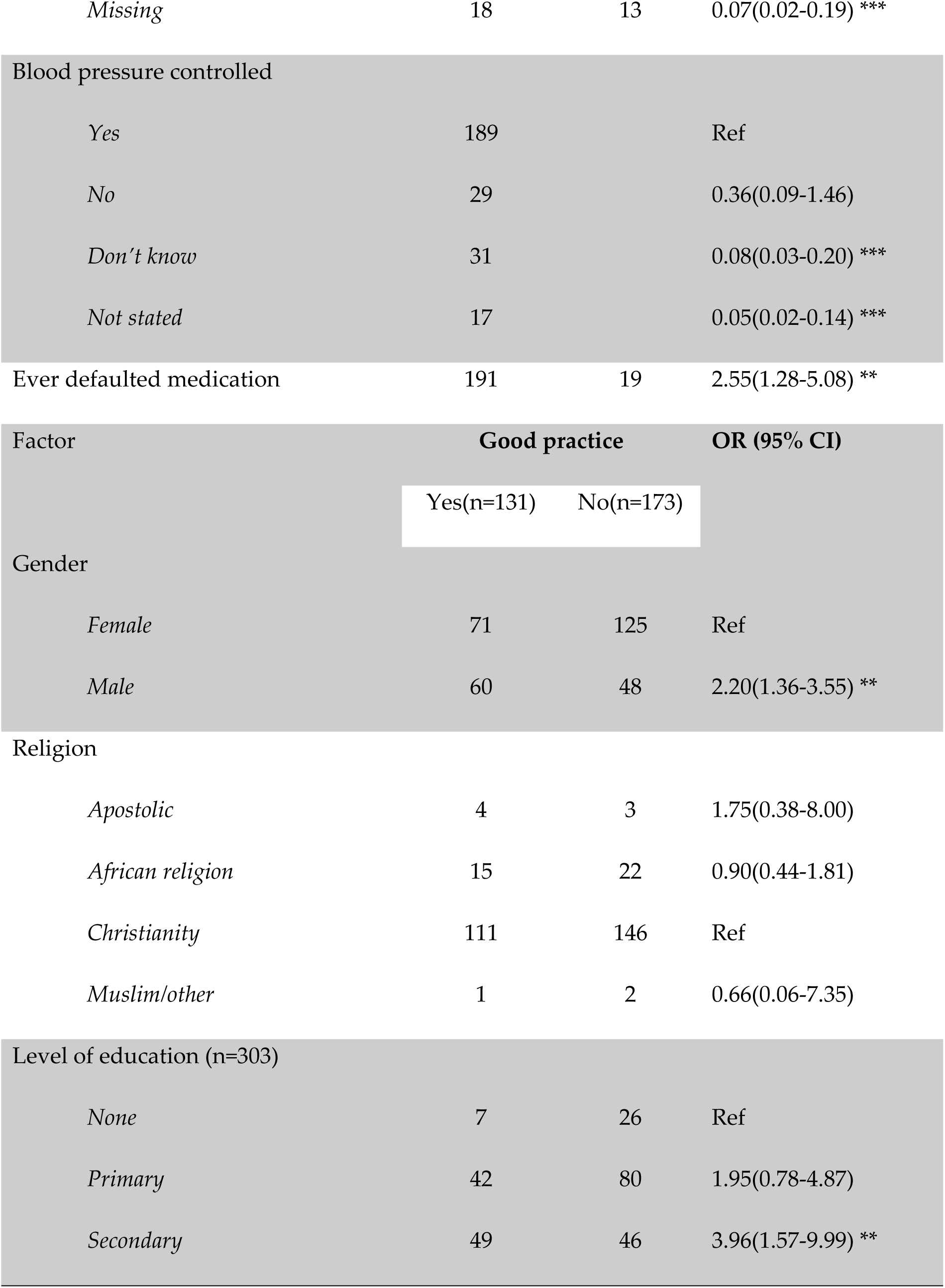
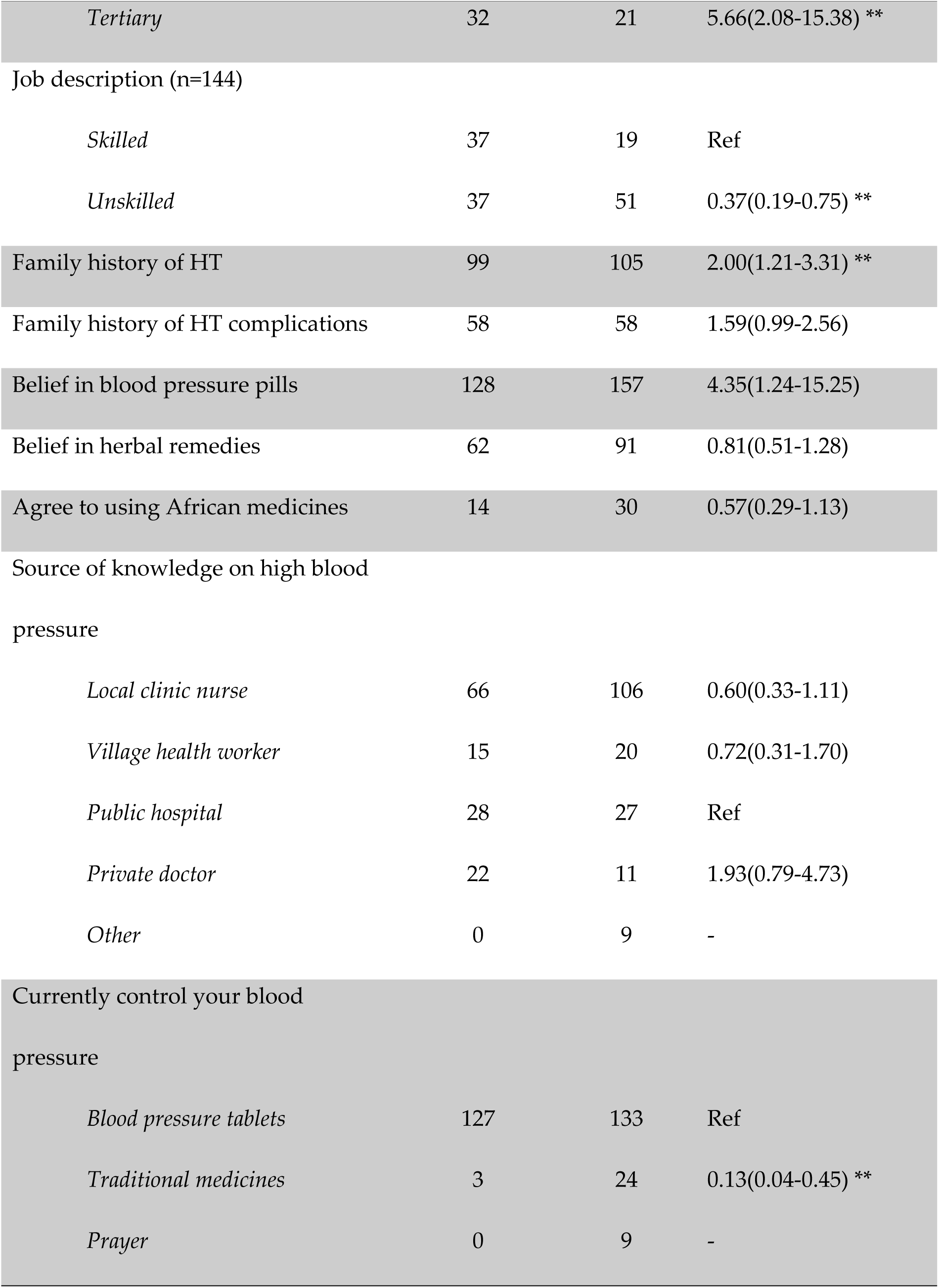
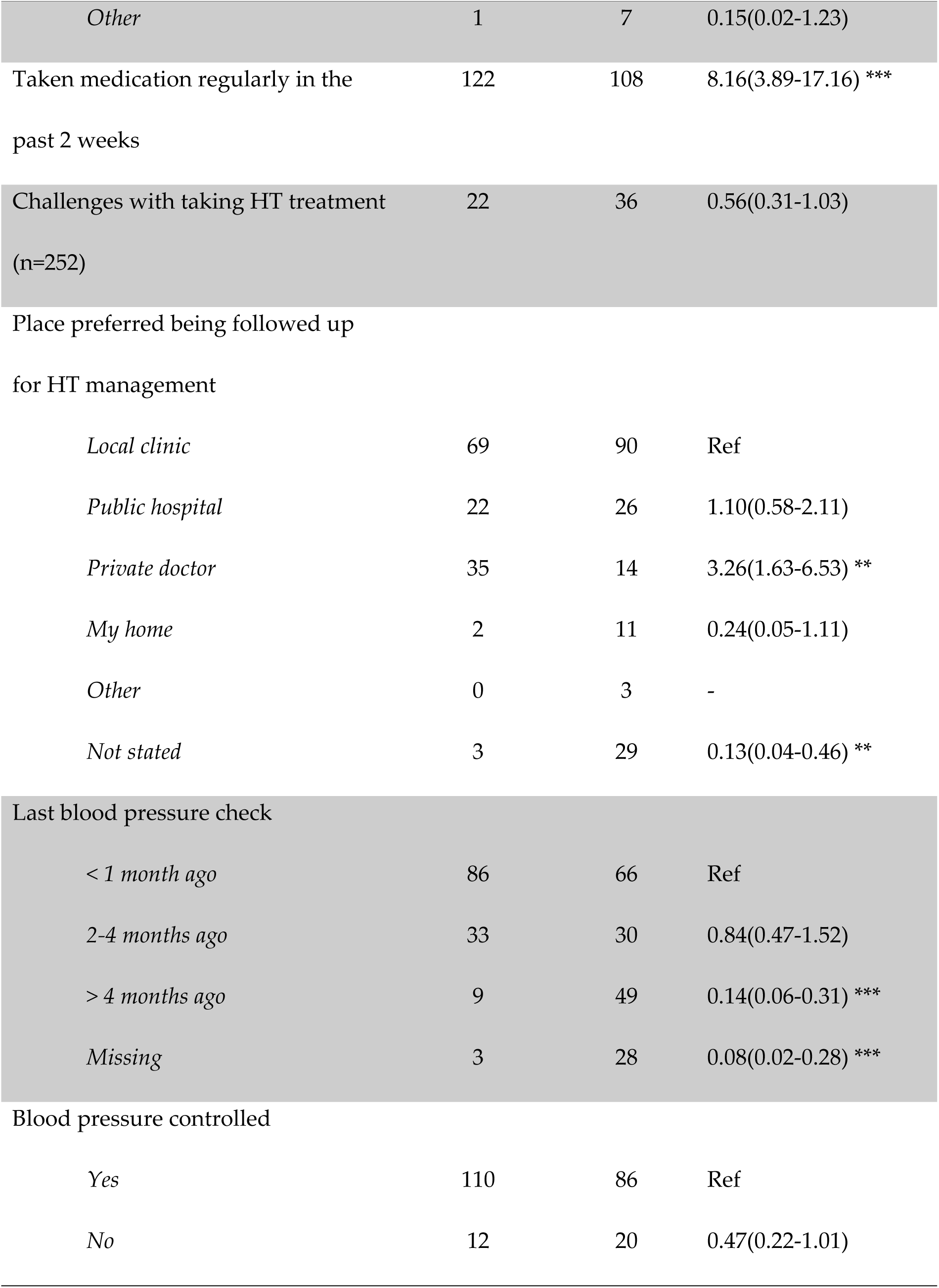
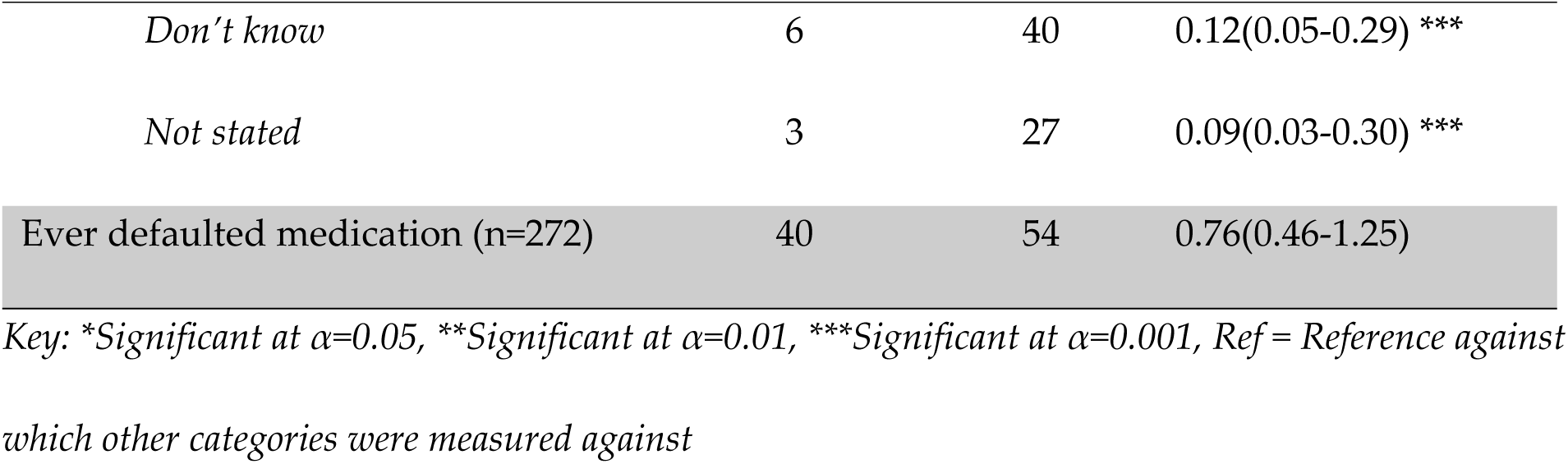
Regression analysis of factors affecting hypertension knowledge in ward 14, Gwanda district community-based action research project on hypertension, May 2017.

In relation to knowledge, those who attained tertiary education and secondary education were 7.52 (95%CI:2.76-20.46) and 3.68 (95%CI;1.61-8.41) more likely to have better knowledge than those who had no formal education respectively. Those that believed in herbal medicines and those that used African traditional medicines were 53% (95%CI:0.29-0.76) and 68% (95%CI:0.29-0.76) less likely to have good knowledge compared to those who did not believe and use traditional medicines respectively.

As far as attitude and practice are concerned, taking Christianity as the standard, those that believe in African religion were 88% (95%CI: 0.05-0.26); less likely to have their blood pressure controlled. There were benefits of education and attitude improved with level of education. Attaining secondary education and tertiary education is 2.86 (95%CI: 1.06-7.71) and 4.59 (95%CI: 1.28-16.44) more likely to have a positive attitude towards hypertension as compared to respondents that have not received formal education. Those that added salt on the table were 60% (95%CI: 0.18-0.88) less likely to have their blood pressure controlled compared to those that did not. Those who believed in the use of blood pressure tablets for controlling HT were 30.45 (95% CI: 10.10-91.79) more likely to have their blood pressure controlled as compared to those who did not while those who controlled their blood pressure with traditional medicines were 97% (95%CI: 0.01-0.08) less likely to have their blood pressure controlled.

Those who confessed having challenges with taking hypertensive treatment were 82% (95%CI: 0.07-0.48) less likely to have their blood pressure controlled. With regards to practice, those with a family history of hypertension were 2.00 (95%CI: 1.21-3.31) more likely to have their blood pressure controlled while those that took medication regularly in the preceding two weeks were (95%CI: 3.89-17.16) more likely to have their blood pressure controlled. There were no statistically significant differences in knowledge across gender, religion and whether respondents were skilled or not in their jobs.

## Discussion

The majority (65%) of PLWHT in this community were females as compared to the men. Some studies reported that hypertension was noted to be prevalent more commonly in females linked with unhealthy diets in low income countries [14]. These findings are similar to most studies on HT awareness treatment and control where women participants were more than men, however both genders are represented [21]. More than half of the participants (51%) were not educated beyond primary school and 11% had no formal education at all. In a community where formal education is low and the persons afflicted by disease are vulnerable due to socio-economic factors, poor health seeking behaviors are common. Educational attainment was directly proportional to knowledge on hypertension as those with tertiary education had better knowledge as compared to those without formal education. Thus, those diagnosed may not have access to treatment and may not be able to successfully control their illness over the long term due to poverty.

This study was conducted in a rural disadvantaged community where supposedly there is little money to spend on health. Basing on observations and declaration of income, most of the respondents were poor and out of pocket health financing was a challenge. Low family incomes could also explain the reduced pattern for abuse of alcohol and tobacco in the community. Similarly, the consumption of fruits was low as they relied mainly on seasonal wild fruits, however vegetable consumption was high on four or more days a week (67%) as they were the commonly available relish. There was an identified risk of consuming too much salt with food (59%) in the community. The Zimbabwe’s National Health Strategy (2009-2013) reported the increase in hypertension prevalence mainly attributed to high salt diet, lack of exercise, tobacco smoking and excess alcohol intake [22].

Value beliefs and practices of an ethnic or racial group within a community can influence acceptance and adherence to health messages as advised by clinicians and academic researchers [19]. The majority of respondents were above 60 years of age and an identified risk factor was family history of hypertension (67%). The majority (94%) believed in using tablets for controlling HT although there are deep beliefs in the use of herbs (51%) and traditional medicines (15%) which influenced their health seeking behavior. We noted that those that had belief in herbs and used traditional medicines had poor knowledge on hypertension and this could contribute to continued myths and misconceptions on hypertension and ultimately poor community outcomes. Although not statistically significant, the older patients were more aware of their hypertensive status; however, studies have shown that this does not universally translate to better hypertension control [21]. This information provides a launchpad for developing community-based hypertension health packages targeted at correcting existing myths, misconceptions and misinformation for improved hypertension management.

There was generally poor knowledge on the risk factors, causes and awareness on hypertension among PLWHT. Most respondents (65%) believed that HT was caused by stress and 17% only knew that the cause is largely unknown. Palpitations were reported as the commonest symptom of HT by 86% of respondents while 7% said it was asymptomatic. It was unanimous among respondents that death was the ultimate outcome of uncontrolled HT. Poverty, ignorance, a poor educational background and weak community health education platforms were determinants of poor knowledge. Significant socio-economic disparities influence the level of HT awareness, treatment and control in LMIC [23]. To compound the knowledge problem in the community, the VHWs were not actively involved in the hypertension care loop.

Hypertension rarely has attributable signs and symptoms in the early stages and many people go undiagnosed [12]. The lack of symptoms for patients with HT contributes to both lack of awareness and reduced compliance to treatment. The improvement in HT control cannot be measurable by symptom relief, thus there is no perceptive benefits for the individual. It is known that hypertension awareness in Zimbabwe is low and this has an impact in low diagnosis, treatment and control hence there is need for a specific policy for prevention and control of hypertension in Zimbabwe [24]. Evidence has shown that even in high income countries, hypertension awareness remains a challenge with 50% of the population being aware of their status and this is estimated to be 40% in LMIC [25-27].

Improved knowledge on hypertension should focus on primary prevention as this is cost effective in low resource settings. Diet (83%) was singled out as the commonest risk factor in developing HT, however 14% had no knowledge of risk factors for HT. Primary prevention reduces the expenses on medical care and the resultant complications of high blood pressure. Awareness screening programs, skills training and capacity building of health workforce on how to deal with hypertension and its associated risk factors including access to low cost antihypertensive medicines are key for developing countries with limited resources [28]. It was noted that 65% of those who took medication perceived that they had well controlled blood pressure however we found out that their scale of measurement was based on experiencing or perceived “complications” rather than blood pressure readings.

In Zimbabwe there are limited national studies on hypertension prevalence while there is lack of infrastructure to enable and support hypertension surveillance [14]. This was evident in that more than 30% of respondents had last checked their blood pressure for more than 4 months while some had lost track of when they had a BP checked. The local clinic was the only place where a blood pressure machine was found however, sometimes the services would be unavailable to various logistical reasons. This then calls for concerted efforts to prioritize service delivery, and funding for HT consumables. Special priority and focus should be on the crafting policy and research-based implementation of tailor-made service delivery packages to reduce hypertension related morbidity and mortality [21].

Nurses were pivotal as a source of HT health (57%) while only 12% reported that they would approach VHWs. A high defaulting rate among of 31% was possibly due to recurrent stock-outs of antihypertensive medicines at the local clinic. Hypertension affects populations negatively in low- and middle-income countries where health systems are weak [11,12,29]. Shortages of medication coupled with long travelling distances to the health facility contributed to poor hypertension outcomes; these findings are reported in other studies as well [21,30,31]. These challenges needed be addressed through primary prevention health education strategies on; treatment compliance, side effects and HT complications while making use of VHWs in community HT care. Several studies indicate that most Africans pay out of pocket for their health bills and these are supplemented by free services subsidized by donors and local governments. However, most of these resources are channeled towards communicable diseases leaving NCDs with little funding [32].

The study had several limitations as there is no standardized instrument to measure HT knowledge, attitudes and practices. We therefore used literature, community knowledge and field experiences to design our data collection tools which may not have been exhaustive. The algorithm used for data collection left some chance of missing hypertensive patients or enrolling patients who may not have been diagnosed of hypertension. It is possible that recruitment bias could have been introduced in that all participants were self-reported hypertensive patients and there was no rigorous verification of hypertension diagnosis.

The study findings were used to identify gaps in knowledge, attitudes and practices including myths and misconceptions by hypertensive patients, village health workers and the community at large on hypertension. These were then used to develop the methodology for the implementation phase of the community participatory action research (CBPR) study we conducted (these study findings were published in a separate paper) [33]. The CIG validated the study findings, and they then used these during the action reflection cycles for planning and learning purposes. Subsequently, the implementation of new community strategies for improved primary prevention of hypertension in the CBPR study [33], were informed by the findings in this baseline quantitative study. Thus, by implication recommendations of improved service delivery from the hypertension CBPR study were influenced partly by the findings in this study.

## Conclusion

Poverty related socio-economic determinants were associated with poor knowledge on hypertension. Health services factors such as medication shortages at the rural health center, poor funding and weak health education platforms were noted in the community. These contributed to reduced awareness and control of hypertension in the community. Concerted efforts were needed to deliberately create community hypertension awareness utilizing community members. Building health worker infrastructure and capacity on hypertension care and an enabling environment for improved disease surveillance for primary prevention of hypertension will benefit disadvantaged communities.

## Acknowledgements

The authors acknowledge the CIG members who participated in this study by adapting it to the community needs of Ward 14 in Gwanda District, Matebeleland South, Zimbabwe. We also acknowledge hypertensive patients who participated in Phase 1 and all the community members of Ward 14 and Vasco Chikwasha the biostatistician who did in-depth data analysis.

## Authors’ contributions

Pugie Tawanda Chimberengwa and Mergan Naidoo (University of Kwa Zulu-Natal) were responsible for conceptualizing the paper. Pugie Tawanda Chimberengwa collected data and wrote the initial manuscript. Mergan Naidoo supervised the study, reviewed and edited the manuscript.

